# Musicians and non-musicians’ consonant/dissonant perception investigated by EEG and fMRI

**DOI:** 10.1101/2021.08.15.456377

**Authors:** HanShin Jo, Tsung-Hao Hsieh, Wei-Che Chien, Fu-Zen Shaw, Sheng-Fu Liang, Chun-Chia Kung

**Affiliations:** Institute of Medical Informatics, National Cheng Kung University (NCKU), Tainan, Taiwan, 70101; Department of Computer Science and Information Engineering, NCKU, Tainan, Taiwan, 70101; Department of Psychology, NCKU, Tainan, Taiwan, 70101; Mind Research and Imaging Center, NCKU, Tainan, Taiwan, 70101

## Abstract

The perception of two (or more) simultaneous musical notes, depending on their pitch interval(s), could be broadly categorized as consonant or dissonant. Previous studies have suggested that musicians and non-musicians adopt different strategies when discerning music intervals: the frequency ratio (perfect fifth or tritone) for the former, and frequency differences (e.g., roughness vs. non-roughness) for the latter. To extend and replicate this previous finding, in this follow-up study we reran the ElectroEncephaloGraphy (EEG) experiment, and separately collected functional magnetic resonance imaging (fMRI) data of the same protocol. The behavioral results replicated our previous findings that musicians used pitch intervals and nonmusicians roughness for consonant judgments. And the ERP amplitude differences between groups in both frequency ratio and frequency differences were primarily around N1 and P2 periods along the midline channels. The fMRI results, with the joint analyses by univariate, multivariate, and connectivity approaches, further reinforce the involvement of midline and related-brain regions in consonant/dissonance judgments. Additional representational similarity analysis (or RSA), and the final spatio-temporal searchlight RSA (or ss-RSA), jointly combined the fMRI-EEG into the same representational space, providing final support on the neural substrates of neurophysiological signatures. Together, these analyses not just exemplify the importance of replication, that musicians rely more on top-down knowledge for consonance/dissonance perception; but also demonstrate the advantages of multiple analyses in constraining the findings from both EEG and fMRI.

**Significance Statement:** In this study, the neural correlates of consonant and dissonant perception has been revisited with both EEG and fMRI. Behavioral results of the current study well replicated the pattern of our earlier work (Kung et al., 2014), and the ERP results, though showing that both musicians and nonmusicians processed rough vs. non-rough notes similarly, still supported the top-down modulation in musicians likely through long-term practice. The fMRI results, combining univariate (GLM contrast and functional connectivity) and multivariate (MVPA searchlight and RSA on voxel-, connectivity-, and spatio-temporal RSA searchlight-level) analyses, commonly speak to lateralized and midline regions, at different time windows, as the core brain networks that underpin both musicians’ and nonmusicians’ consonant/dissonant perceptions.

## Introduction

In Western tonal music, an octave is defined by 12 equal spaced semitones, whose distance in turn defines the pitch interval. For example, “Do” and “Mi” are 4 semitones, or two full notes, apart (Aldwell et al., 2010). Playing 2 or more pitch intervals together, or so-called a chord or harmony, is the structural foundation of melody (Krumhansl, 2001). Intervals could be broadly categorized as consonant and dissonant: the consonant intervals are usually considered as stable, pleasant, and more acceptable (Malmberg, 1918); whereas the dissonant ones as more unstable, unpleasant, and tension-building (Rameau, 2012). Perceptual differences between consonant and dissonant intervals are found across different ages (Bidelman & Heinz, 2011), across various species, such as primates ((Perani et al., 2010; Virtala et al., 2013) and chicks (Chiandetti & Vallortigara, 2011), and regardless of musical backgrounds (Koelsch, 2011; Peretz, 2002). On the other hand, dissonance has been related to another characteristic called ‘roughness’ (Vassilakis & Kendall, 2010), when two close-by frequency intervals interfere with each other, generating ‘beating’ vibrations (Daniel, 2008; Zwicker et al., 1957), as stated by Helmholtz’s psychoacoustic theory (Lichtenwanger et al., 1954)

There have been quite a number of neuroimaging studies addressing the consonance and dissonance perception of musicians (and non-musicians). They include: (a) Event-Related Potential (ERP) studies (Kung et al., 2014; Minati et al., 2009; Regnault et al., 2001), that commonly identify the N1 (~100ms) and P2 (~200ms) components along the midline electrodes; (b) fMRI studies on the associated neural substrates (Bidelman & Krishnan, 2009; Leipold et al., 2021): the primary auditory cortex and superior temporal gyrus (STG), that are jointly implicated in pitch discriminations, rhythm judgments, and higher order music appreciations (Peretz & Zatorre, 2005); (c) Magnetoencephalography (MEG) studies (Kuriki et al., 2006; Tabas et al., 2019), that provide evidence of early (63 ms) processing of auditory cortex in both musicians and controls; and even the (d) electrocorticographic (ECoG) study (Foo et al., 2016) demonstrating high gamma (~70-150Hz) in the right superior temporal gyrus, or r-STG, of epilepsy patients (non-musicians), especially in dissonance condition. Together, these methodologies provide both the diversive, owing to each method’s sensitivity niche in the spatiotemporal axes, and combinatory view into the training-induced neuroplasticity of musicians’ (vs. non-musicians’) brain. However, these pursuits not surprisingly lack the consistency in the design protocol, stimuli, and the musicians’ degree of expertise, etc; rendering summaries of these findings taken at the abstract level, at best.

Recently, there is a growing concern for the reproducibility of neuroscientific, here fMRI, findings (Botvinik-Nezer et al., 2020; Tom et al., 2007), partly originated from the earlier seminal work (Camerer et al., 2018; Nosek et al., 2015). To our knowledge, to date very few, if none, replication studies to reassure the findings of related works in musicians’ consonant/dissonant perception (Merrett et al., 2013).

The purpose of the present study is twofold. First, to extend our previous research (Kung et al., 2014) by first replicating the behavioral and neurophysiological findings in another group of musicians and non-musicians. Second, the same ERP experimental protocol will be applied to the fMRI setting, with minimal but necessary adjustments, to investigate the neural correlates of similar behavioral performance (or separate reliances on different musical features by musicians and non-musicians). The focus will be equally emphasized on both the checking of replication of the behavioral and ERP findings, and subsequently the separate fMRI analyses by multiple approaches, including univariate (aka. General Linear Model, or GLM), multivariate (aka. Multi-Voxel Pattern Analysis, or MVPA, searchlight) (Kriegeskorte et al., 2006), and connectivity (including psychophysiological interactive, or PPI (O’Reilly et al., 2012); as well as representational similarity analysis, or RSA (Nili et al., 2014) and ssRSA ((Salmela et al., 2018; Su et al., 2014)), results. Lastly, the joint analyses on the ERP and fMRI data would provide the final converging evidence of the common neural substrate of asserted neural signatures.

## Materials and Methods

### Participants

The musician subjects were recruited from the department of music of the National University of Tainan. Among the 40 amateur musicians who participated in the behavior dissonance/consonance judgement tasks (for screening), 15 were above-75% accuracy performance and invited to participate in ERP and fMRI experiments. Selected musician participants were all female with varied types of musical instruments background. The average age of musicians was 21.7±1.3 yrs old, with 14±4.3 years of average musical training background, and 8.4±5.30 weekly training hours. For the non-musician group, 30 NCKU students with no formal musical training nor musical instrument experiences were recruited for behavior experiments. There are 14 subjects for the ERP, and 17 for the fMRI experiments, respectively. Their average age was 21.9±1.7 yrs old.

### Stimuli and Procedure

The same stimuli from Kung, et al. (2014) were adopted: two types of pitch interval: tritone (6 semitones, dissonant intervals) and perfect fifth (7 semitones, consonant intervals) were generated by mixing two sinusoidal tones. Each type of pitch intervals contained 10 dyads tuned to the equal-tempered chromatic scale (A4 = 440 Hz), and the dyads were evenly distributed within the defined frequency range of G#2 (104HZ) to Eb5 (622HZ). In addition, each sound stimulus was orthogonally manipulated by frequency differences, coded as ‘roughness’ and ‘non-roughness’, based on the curve of the critical band width (Zwicker et al., 1957). The 4 categories: “tritone with roughness” (T1-T5), “tritone without roughness” (T6-T10), “perfect fifth with roughness” (P1-P5), and “perfect fifth without roughness” (P6-P10) were also the same arrangements as in Kung et al. (2014).

The behavioural experiments were administered prior to EEG and fMRI experiments. The procedure of behavioral and ERP experiments have been detailed in Kung et al., (2014, pp. 973), with a few highlights here: for the behavior responses, subjects were instructed to listen attentively to each stimulus presentation, judged the relative consonance, and pressed a response button as quickly and accurately as possible following the stimulus onset. No feedback was given on their response. The stimulus sets consist of 20 two-note intervals, randomly presented in two consecutive sessions (40 trials in total, each stimulus presented 2 times). The musicians with above 75% accuracy were further recruited for the neuroimaging experiments.

In the subsequent ERP experiment, invited participants were told to silently attend, but not behaviorally respond with button presses, to the presented stimuli, thereby avoiding contamination of associated cortical responses (since CZ and FZ electrodes are close to the motor cortex). The inter-stimulus interval (ISI) was multiples of 4.5s (e.g. 4.5s, 9s, or 13.5s, etc), thereby making the flow of ERP trials similar to those in the later fMRI counterpart. The ERP experiment contained two 15-minute sessions, each with 7 repetitions of 20 trials that were randomly presented, with 5 minutes break in between sessions. Plus the ~30 minutes of preparation, each ERP experiment took about 70 minutes to complete.

For the fMRI experiments, an extra concern about the scanner noise interfering with the audio presentations was addressed by adopting the ‘sparse sampling’ design (González-García et al., 2016; Hall et al., 1999; Perrachione & Ghosh, 2013; Whitehead & Armony, 2018), where the baseline/actual scan was separated from the stimulus presentation, with the necessary silent gap in between for better hearing/evaluation (See Fig. 1 for details). The 4.5s TR (2s scan+ 1s pre-sound gap+ 0.3s sound + 1.15s after-sound response period) was determined after several rounds of pilot experiments, calibrating the peri-audio gap durations (from .5s to 1s), finding the best timing arrangements so that most subjects can hear the sound stimuli clearly, and do the subsequent consonant/dissonant judgment as accurately as in the behavioral sessions (reported in the results section). Even so, for each participant, the experimenters still have to adjust the volume of the NNL ear-plugged headphone (http://fmri.ncku.edu.tw/tw/equipment_2.php), so that each participant can hear clearly and comfortably before 6-10 runs of formal fMRI experiment (each run containing 2 rounds of 20 trials). For each finished participant, the sum of subject payment was NT$1000 (~$33 USD).

**Fig. 1.**
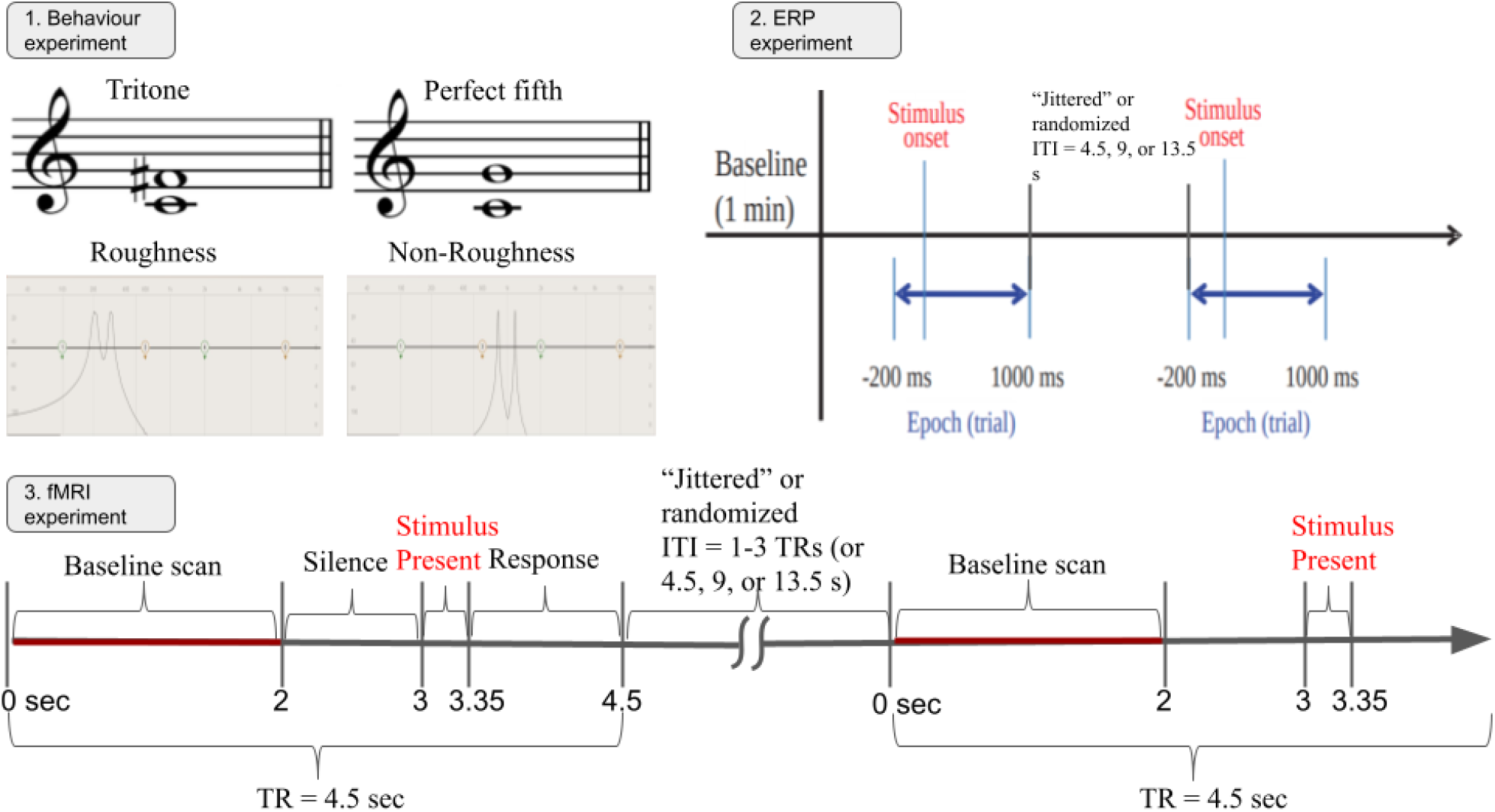
The illustration of the (1) behavioral, (2) ERP, and (3) fMRI experimental design, each carried out in separate sessions. First, the two-tone intervals were divided into 4 categories: tritone, perfect fifth, rough, and non-rough, along the frequency ratio (tritone vs. perfect fifth) and difference (rough vs. non-rough) dimensions. The sheet notes in the upper two columns: tritones and perfect fifth, represent the idea of interval ratio along two pure tones (as was done in the current experiments), not along the two pitches. The behaviour experiment was to test the expertise level in consonance/dissonance judgment task for both the musician and non-musician participants. Musicians have to be above 75% performance for further recruitment into ERP and fMRI studies, whereas non-musicians were invited based on their willingness and availability (not by task performance). In the ERP experiment, subjects were told not to respond behaviorally. In the fMRI experiments, the sound stimuli were presented in between the two silence periods (1s and 1.15s) for clearer hearing, and the effective baseline scan only lasted for 2s. The jittering was implemented by the 1-to-3 times of TR (4.5s) as the inter-trial-interval (ITI), or 4.5, 9, or 13.5 s. To equate the ERP and fMRI experiments, the inter-trial intervals were kept identical (1-3 multiples of 4.5s TR) across the two protocols.

### Behavioural data analysis

Each subject’s behaviour responses of the dissonance/consonance judgment task, mainly the average accuracy and response time (RT), were calculated to show the differences between musicians and non-musicians. The group and conditional interaction effects entered the analysis of variance (ANOVA), to reveal the response differences in perception judgment across frequency intervals (perfect fifth vs. tritone) and frequency differences (rough vs. non-rough).

### EEG Acquisition, Preprocessing and ERP Analysis

Due to the fact that the EEG/ERP experiment was replication in nature, the ERP preprocessing and analysis procedures will be mostly the same as described in Kung et al., (2014, pp. 973-974,, with minor additions): scalp EEG was continuously recorded in DC mode at a sampling rate of 1000HZ from 32 electrodes mounted in an elastic cap and referenced to the mastoid bone, in accordance with the 10-20 international system. Impedance at each scalp electrode was reduced to 5KΩ or below. Electrode positions, physical landmarks, and head shape were digitized using Polhemus Fastrak digitizer and 3D spaceDx software contained within the Neuroscan SCAN software package. The EEG was amplified using Neuroscan NuAmp and filtered on-line with a bandpass filter (DC to 100Hz, 6-dB/octave attenuation). The EEG data were down-sampled (256 Hz)and filtered with a band-pass filter 0.5-30 Hz (EEGLAB, FIR filter) (Delorme & Makeig, 2004) to eliminate slow drifts as well as muscular artifacts. The data from 200 ms prior to and 1000 ms after the onset of each stimulus were segmented and baseline-corrected by the pre-stimulus period average. Each epoch exceeding over 100 in amplitude was automatically rejected at any EEG electrode, remaining accepted trials are averaged by condition types.

For the analysis of ERP data, the 8 peri-midline (F3, F4, FC3, FC4, C3, C4, CP3, and CP4) and 4 midline channels (FZ, FCZ, CZ, and CPZ) were our primary focus. Earlier EEG studies indicated that the midline channels play an important role in consonance judgments for musicians (Itoh et al., 2003, 2010; Maslennikova et al.,2015). The group differences between musicians and non-musicians were therefore tested by ANOVA across the above mentioned 4 channels to see at which components were significantly different, and 12 channels (4 midline channels + 8 peri-midline channels) were individually analyzed to identify significant contrasts for each group (e.g., tritone vs. perfect fifth; rough vs. non-rough).

### fMRI Acquisition, Preprocessing and GLM Analysis

fMRI images were acquired with a 3T General Electric 750 MRI scanner (GE Medical Systems, Waukesha, WI) at the NCKU MRI center, with a standard 8-channel array coil. Whole-brain functional scans were acquired with a T2* EPI (TR= 2 s, TE= 35 ms, flip angle = 76 degree, 40 axial slices, voxel size = 3 × 3 × 3 mm). High-resolution structural scans were acquired using a T1-weighted spoiled grass (SPGR) sequence (TR=7.5s, TE=7.7ms, 166 sagittal slices, voxel size = 1 × 1 × 1 mm). The fMRI data were pre-processed and analyzed using BrainVoyagerQX v. 2.6 (Brain Innovation, Maastricht, The Netherlands) and NeuroElf v1.1 (http://neuroelf.net). After skipping the first 5 prescan volumes (default option), the first 2-3 functional volumes were again discarded (due to residual T1 saturation effect), followed by slice timing correction and head motion corrections (with six-parameter rigid transformations, and aligning all functional volumes to the first volume of the first run). Neither high-pass temporal filtering nor spatial smoothing was applied. The resulting functional data were co-registered to the anatomical scan via initial alignment (IA) and final alignment (FA), and then both fmr and vmr files were transformed into Talairach space.

The Volume Time Course (VTC) files were created with corresponding Stimulation Design Matrix (or SDM file), and entered the general linear model (GLM) with random-effect (RFX) analysis. Additionally, Psycho-Physiological Interaction (PPI) analysis (O’Reilly et al., 2012) was conducted to show brain regions functionally connected to the seed region, defined from the GLM contrast (e.g., tritone > perfect fifth from the musician group). All the results were cluster-thresholded with FWE correction using Alphasim at p < 0.05 (corrected), and visualized with 4 surface renderings (e.g., medial and lateral views for both LH and RH).

### Multivariate analysis

For multivariate analysis, MVPA searchlight mapping (Kriegeskorte et al., 2006) was applied by estimating each trial’s beta weights with Least Square Separate (LSS) methods (Mumford et al., 2012). The resulting estimation was z-normalized and demeaned prior to entering the CoSMoMVPA (Oosterhof et al., 2016) under MATLAB R2018a for MVPA searchlight analysis (default cluster 100 voxels for each tested voxel). The Linear Discriminant Analysis (LDA) classifier was adopted to train and classify the tritone/perfect_fifth categories in musicians, and rough/nonrough categories for non-musicians, in a leave-one-run-out cross validation manner. The resulting mean cross-subject classification accuracy whole-brain results, each voxel against 50% chance performance, was t-tested for group inferences.

Second, RSA was used in three different ways. The first is the (traditional) voxel-based RSA, which correlates the constructed model RDMs with the fMRI RDMs in a whole-brain searchlight manner (Diedrichsen & Kriegeskorte, 2017). The procedure was to construct eight RDM models (the first four were based on the stimulus dimensions; the latter four were by subject/group responses. see Figure 5 for details) first: (a) all separate: the four stimulus conditions (tritone-rough, tritone-nonrough, perfect fifth-rough, and perfect fifth-nonrough) were equal distance (or dissimilar) to one another; (b) frequency interval-based (tritone vs. perfect fifth) equal dissimilarity; (c) frequency difference-based (roughness vs. non-roughness) equal dissimilarity; (d) high - low frequency gradients: subtracting the means of each two-note intervals across all the twenty stimuli, rendering the frequency-gradients from below- to above-zero; (e) behavioral consonance judgments for the twenty stimuli for musicians; (f) behavioral consonance judgments for the twenty stimuli for non-musicians; (g) behavioral RTs of the consonance judgments by musicians; and (h) behavioral RTs of the consonance judgments by non-musicians. All 8 RDM models were rank-transformed and scaled into 0-to-1 dimension for RSA analyses, to identify the brain regions that were highly associated (with subject random effects sign-rank tests with FDR correction) (Nili et al.,2014).The individual RSA graph was against null (zero) correlation, and group-wise t-tests were applied to yield the final brain maps, overlaying to the BrainNet Viewer (Xia et al., 2013)for comparison with the network-based RSAs.

The second RSA analysis was ROI- and network-based, in which the 90 nodes (from the Automated Anatomical Labeling 3 (AAL3) atlas, from which we selected only the 90 cerebral regions out of the 116 ROIs) (Rolls et al., 2020) were first compared with the 6 (per subject group) model RDMs, to identify the significant ROIs that showed high similarities. The ROI clusters above the threshold for each model were extracted for pairwise correlation to reveal the connectivity between ROI RDMs, later the statistical significance in p-value were converted to z score to scale the size of node with the location of center of gravity in each ROIs from AAL 90. The connectivities among surviving ROIs were computed as well, thereby generating the statistically significant edges which, along with the nearby nodes, formed the network-based graphic representations (Fair et al., 2009). To be concise, the 1st and the 2nd RSA analysis results were overlaid together in Fig. 6. The dissimilarity, or correlation distance measures, were carried out by subtracting correlation values (between beta values and model RDMs)from 1. And then the ROI- or connectivity-based dissimilarity values were transformed into groupwise t-values.

The third, and the last, was the spatio-temporal searchlight RSA (or ss-RSA), which correlates both EEG and fMRI data by their temporal and spatial RDM (Salmela et al., 2018; Su et al., 2014). The average ERP of the 4 midline channels (e.g., Fz, FCz, Cz, CPz) was extracted for each stimulus, and used to construct RDMs across the 307 timepoints, from 200ms prior to 1000ms (with 3ms time bin) after the onset of stimuli, in the EEGLAB (Delorme & Makeig, 2004); and then further correlated with the spatial RDM constructed from estimated fMRI whole-brain betas for each stimuli. The spatio-temporal searchlight RSA was performed between 0ms to 500ms after stimulus onset (see supplementary video 1 and 2 for details). The CoSMoMVPA toolbox (Oosterhof et al., 2016) was used for preprocessing and the searchlight RSA. For the statistical inference, the results of correlation r values were transformed into z scores, before entering into the t-test for RFX group analysis.

## Results

### Behaviour results

The behaviour responses were separately collected for both behavioural and fMRI experiments (no behavioral responses for ERP). In Fig. 2, the distribution of dissonance/consonance judgement ratio across 20 sound stimuli are shown. The average accuracy of musicians’ judgements were 76% (sd = 14.0%), and nonmusician’s judgement 61% (sd = 10.4%), and the average RT for musicians’ responses were 1613.8 ms (sd = 148.6 ms), non-musicians 1347.8 ms (sd = 67.8 ms). The repeated-measure ANOVAs on accuracy across the 4 categories were all significant between musicians and non-musicians: for tritone (F_(9,540)_ = 6.1047, *p* = 3.39 × 10^-8^), perfect fifth (F_(9,540)_ = 5.1556, *p* = 1.00 × 10^-6^), roughness (F_(9,540)_ = 6.2091, *p* =2.33 × 10^-8^) and non-roughness (F_(9,540)_ = 4.2829, *p* = 2.16 × 10^-5^). The overall between-group (32 musicians vs. 30 non-musicians) accuracy difference across all 20 stimuli and 2 sessions was t_(122)_ = 6.3955, *p* = 3.07 × 10^-9^. The overall group difference in response time was not significant (mean t_(122)_=1.7319, *p* > 0.05), except for the stimuli T3 (t_(122)_=2.2498, *p* < 0.05), out of 20 tests.

**Fig 2.**
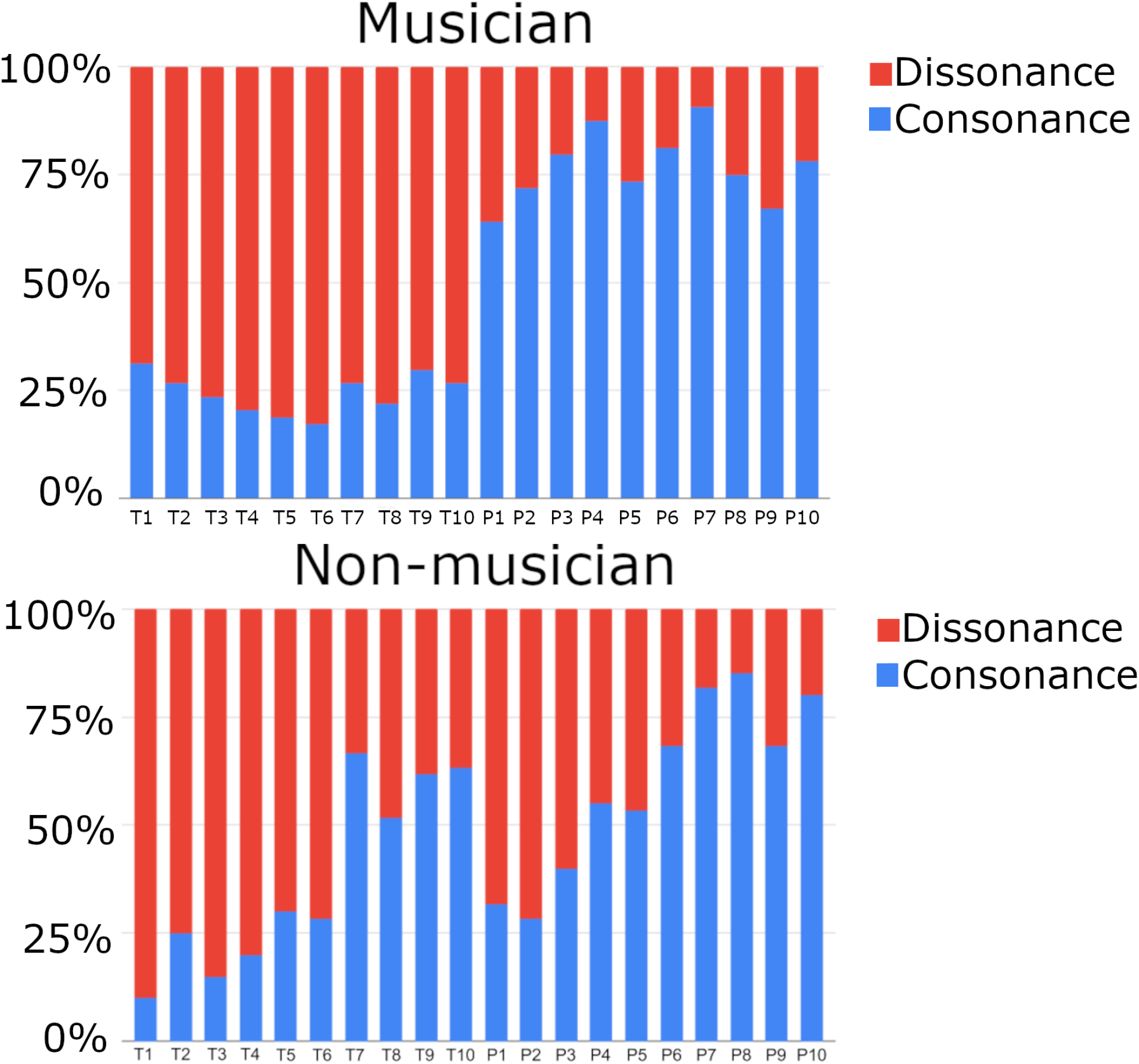
The behavioural responses across four category discrements (tritone, perfect fifth, orthogonalized with rough and non-rough conditions) between the two groups (Bar graph on top for musicians, and the bottom for non-musicians). T1 ~ T5 represent tritone with roughness, T6 ~ T10 tritone without roughness (or non-roughness); P1 ~ P5 perfect fifth with roughness, and lastly P6 ~ P10 perfect fifth without roughness. The percentage of judgments as dissonance were described in red, and the judgment as consonance described in blue color bars.

Not surprisingly, both musicians and non-musicians showed high correlations in accuracy ratio across the stimuli between behavior and fMRI sessions (musician: r_(18)_ = .71, *p*= 4.00 × 10^-4^; non-musician: r_(18)_ = .94, *p*= 3.82 × 10^-10^). As shown in Fig. 2, two groups’ consonance judgment performances matched the predictions made by the “musicians: tritone vs. perfect fifth” and “non-musicians: rough vs. non-rough” dimensions. Importantly, the behaviour responses well replicated our previous results using the same 20 stimuli (c.f., Fig. 4, Kung et al., 2014, pp. 975). In addition, the worry about, and the careful fMRI procedure for each musician subject to avoid, the potential contamination of scanner noise interfering with the auditory stimuli, was cleared and addressed by the positive correlation of 13 early/available musician’s performance both inside and outside the MRI scanner (r_(11)_= .56, *p* < 0.05). Although the stimuli, testing environment, and some of the experimenters were identical between our previous work (Kung et al., 2014) and the current study, the new group of participants, both musicians and non-musicians alike, still performed closely matched, speaking to the reliability of behavioral performance, the most important predecessor to all the underlying brain mechanisms.

**Fig 3.**
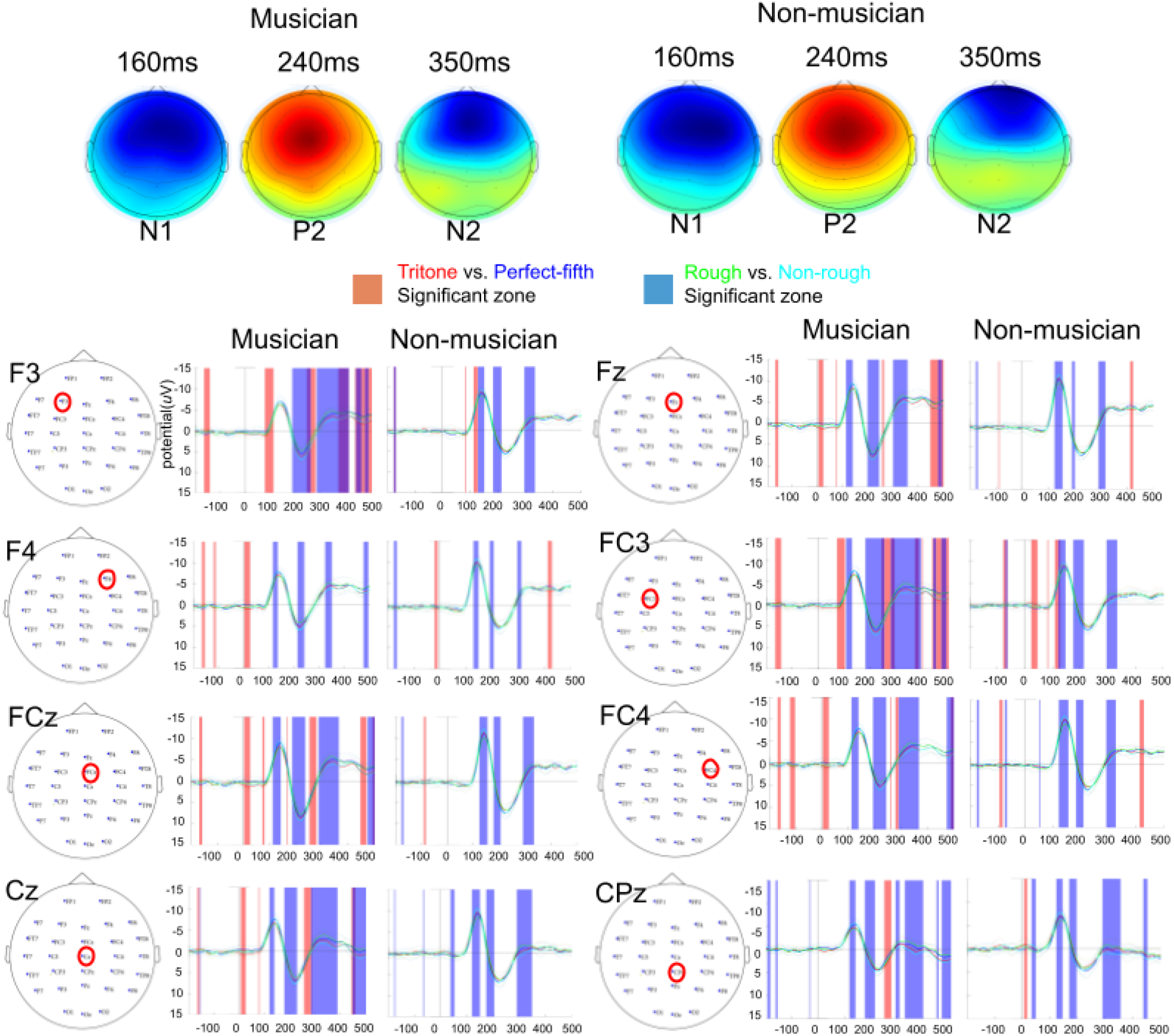
Top row: the topographic plots shown at 160, 240, and 350 ms, for both musician and non-musician groups; and the 2nd to the 5th rows: channel-wise EEG comparisons across F3, F4, FCz, Cz (left column), and Fz, FC3, FC4 and CPz (right column) channels with between musician (left) and non-musician (right) groupwise average time courses differences. Color line matchings: red: tritone, blue: perfect fifth, green: rough, and cyan: non-rough. Orange and blue patches represent the time periods where the two (orange for red/tritone vs. blue/perfect fifth; and blue for green/rough vs. cyan/nonrough, respectively) conditions were significantly different (uncor. *p*= .05).

**Fig. 4.**
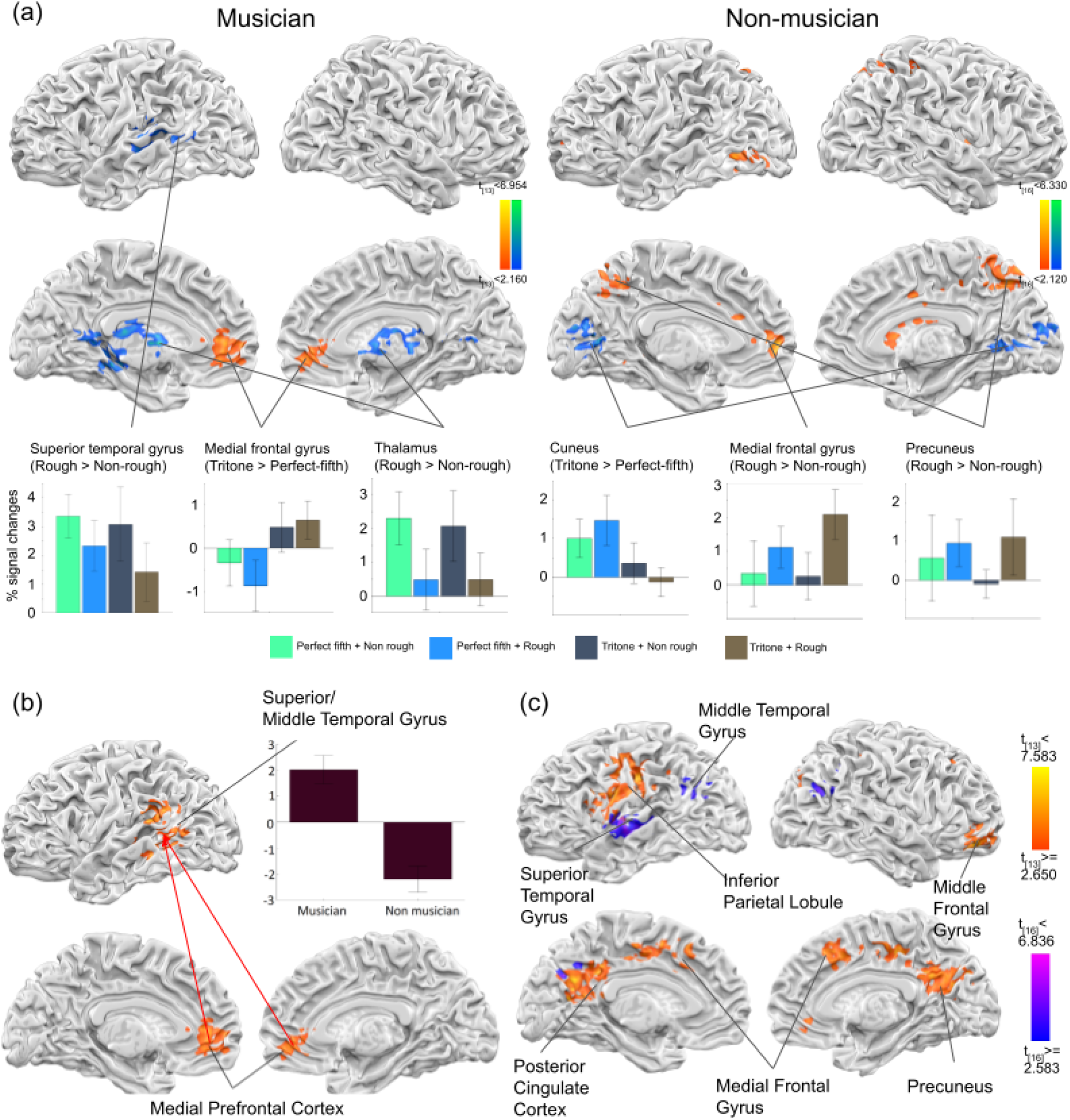
(a) Groupwise GLM results. Both orange and blue colored regions represent the positive and negative group main effects, and for musicians and non-musicians, respectively. For example, the lower left-most bar graph showed higher STG activities for non-rough conditions (relative to those for rough conditions) on musicians; or the 5th (2nd to the rightmost) bar graph represents higher MFG activities for rough conditions than for non-rough conditions, on non-musicians. The bottom beta-plots show the ROIs activation difference across the 4 conditions (from left to right, perfect fifth without roughness, perfect fifth with roughness, tritone without roughness, and tritone with roughness). The GLM results are shown in *p* < .05 with FWE corrected cluster size using alphasim. (b) The seed region, medial prefrontal cortex (below, MPFC, defined from the GLM contrast of tritone > perfect fifth in the musician group) was used for the Psycho-Physiological Interaction (PPI) analysis, and the resulting superior and middle temporal gyrus as functionally connected to MPFC, in the group contrast (musician > non-musician). The bar graph indicates the beta values of conditional contrasts (Tritone > Perfect fifth) × VOI (MPFC) in superior and middle temporal gyrus for each group. (c) MVPA searchlight results of musicians classifying perfect fifth vs. tritone (orange to yellow), and non-musicians between roughness and non-roughness (blue to purple). One can see, once again, the midline + lateralized areas for separate groups, respectively.

### ERP results

In the ERP analysis, one main focus was to identify the electrodes of interests subserving the groupwise consonant judgments, and their associated amplitude differences along the temporal dimension (e.g., N1, P2, etc). For that purpose, the Auditory Evoked Potential (AEP) N1 (150-180 ms), and P2 (180-250 ms)components across the individual electrodes are mainly focused, and put into ANOVA and t-test for comparing conditional differences.

The ANOVA results showed the significant group main effects in the frontal and midline electrodes at the N1 component: F7 (F_(1, 115)_= 9.0307, *p* < 0.005), F3 (F_(1, 115)_=7.1556, *p* < 0.01),F4 (F_(1, 115)_= 10.4919, *p* < 0.005), FT7 (F_(1, 115)_= 10.3878, *p* < 0.005), FCz (F_(1, 115)_= 9.4795, *p* < 0.005), FC4 (F_(1, 115)_= 15.2379, *p* < 1.64 × 10^-4^), Cz (F_(1, 115)_= 13.5431, *p* < 3.64 × 10^-4^), C4 (F_(1, 115)_= 7.3558, *p* < 0.01), and CPz (F_(1, 115)_= 10.1836, *p* < 0.005). At the P2 component, the FC3 (F_(1, 115)_= 7.2016, *p* < 0.01) and FCz (F_(1, 115)_= 7.3397, *p* < 0.01) channels are found to be significant. The interaction effects with group x frequency interval conditions didn’t reveal any significant results in the N1 and P2 components, meanwhile, the interaction effects of group x frequency difference conditions are found at the N1 component in the channel P7 (F_(1, 115)_=5.5301, *p* < 0.05).

In the within group comparison between perfect fifth vs. tritone condition, the musicians showed stronger N1 for tritone then for perfect fifth was found: as early as 160ms, FCZ (t_(14)_= 2.1790, *p* <0.05), CZ (t_(14)_= 2.1564, *p* < 0.05), F3 (t_(14)_= 2.2742, *p* < 0.05), FC3 (t_(14)_= 2.7628, *p* < 0.02), and C3 (t_(14)_= 2.2584, *p* < 0.05). Regarding P2 components, the stronger amplitude for tritone than for perfect fifth was observed as early as 180ms, in the F3(t_(14)_= 2.4215, *p* < 0.05), FZ (t_(14)_= 2.3459, *p* < 0.05), FC3 (t_(14)_= 2.7628, *p* <0.02),FCZ (t_(14)_= 2.5161, *p* < 0.05), C3 (t_(14)_= 2.5231, *p* < 0.05), and CZ (t_(14)_= 2.3779, *p* < 0.05) channels. For the non-musician groups in the contrast of perfect fifth and tritone conditions, the significant results are identified in the CPZ (t_(14)_= −2.1969, *p* < 0.05), CP4 (t_(14)_= −2.3812, *p* < 0.05), and PZ (t_(14)_= −3.1080, *p* < 0.01) channels in the P2 components. In the comparison of amplitude between roughness vs. non-roughness conditions, musicians showed significantly stronger P2 component for roughness than non-roughness conditions in the C3 (t_(14)_= 3.7391, *p* < 0.005), CZ (t_(14)_= 3.3411, *p* < 0.005), CP3 (t_(14)_= 5.0945, *p* < 1.633*10^-4^), CPZ (t_(14)_= 3.6895, *p* < 0.005), and CP4 (t_(14)_= 3.4101, *p* < 0.005) channels as early as 180ms, additionally, FT7 (t_(14)_= 3.6930, *p* < 0.005), F3 (t_(14)_= 3.6881, *p* < 0.005), FC4 (t_(14)_= 3.4735, *p* < 0.005), F3 (t_(14)_= 4.7869, *p* < 2.896*10^-4^), FP1 (t_(14)_= 3.5026, *p* < 0.005), and FP2 (t_(14)_= 3.4146, *p* < 0.005) channels were identified around 200 ~ 220ms. In the nonmusician group, the significant differences in rough vs. nonrough were identified in the N1 component including FZ (t_(14)_= −3.3325, *p* < 0.01), FCZ (t_(14)_= −3.3501, *p* < 0.01), FC4 (t_(14)_= −3.1156, *p* < 0.01), CZ(t_(14)_= −4.5077, *p* < 5.889*10^-4^), C4 (t_(14)_= −3.6895, *p* < 0.01),and CPZ (t_(14)_= −3.5906, *p* < 0.01) channels, and the P2 component only including the C4 (t_(14)_= 3.2482, *p* < 0.01) channel.

To sum up the ERP results, compared to our previous ERP study results (Kung et al., 2014), the current study showed similar findings for musicians along the frequency ratio dimension (“tritone vs. perfect fifth”), especially in FZ and FCZ channels, along the N1 (150-180 ms) and P2 (~250ms) periods. For the frequency differences dimension, however, in the current study both musicians and non-musicians exhibited similar N1 and P2 effects along the midline and frontal electrodes, unlike the previous effect for non-musicians only (cf. Fig. 6, pp. 977, of Kung et al., 2014). These partial replications, more in line in frequency ratio (yes for musicians and no for non-musicians; bot replicated), but less so in frequency differences (both yes for musicians and non-musicians in the current study; unlike ‘no for musicians and yes for non-musicians’ reported in Kung et al.) results, still speak to the reliability of related findings (Itoh et al., 2003), and set a foundation for the later fMRI extensions. More speculations about the ERP differences will be listed in the discussion section.

### fMRI ANOVA-PPI-MVPA searchlight results

The group (musicians vs. non-musicians) x 2 (frequency intervals: tritone vs. perfect fifth) x 2 (frequency differences: rough vs. nonrough) 3-way ANOVA revealed the group main effect in the posterior cingulate cortex (PCC), precuneus, left inferior parietal lobule (l-IPL), inferior frontal gyrus (l-IFG), and caudate (shown in supplementary figure 1 and table 1, orange color). Meanwhile, the group by frequency interval interactions (tritone vs. perfect fifth) identified only the right inferior parietal lobule (r-IPL, shown in yellow), and the group by frequency difference (roughness vs. non-rough) interaction revealed (in pink color): the nucleus accumbens (NAcc), dorsal medial prefrontal cortex (dmPFC), cingulate gyrus, and precuneus. To dig further, the separate group GLM results showed that only musicians have significant activation differences in the bilateral medial prefrontal cortex (MPFC, see Fig. 4a bottom for visualizations) for frequency interval contrast (tritone > perfect fifth), and not so in the non-musician group. Along the frequency difference dimension, both groups exhibited the significant activation differences in the left superior temporal gyrus (l-STG), and thalamus for musicians (higher in the “rough than non-rough” condition); and the MPFC, middle cingulate cortex, and precuneus for non-musicians (higher in the non-rough than the rough condition, opposite to that of the musicians).

For the functional connectivity analysis, out of the several seed regions that were explored, we only found the MPFC seed showing stronger connections in left STG, and Middle Temporal Gyrus (MTG) in the contrast of ‘tritone > perfect fifth’, and more in musicians than non-musicians (see Fig. 4b). Those temporal gyrus were actually close to the Heschl gyrus that was reported in a recent study (Bravo et al., 2020) that employed the similar PPI analysis with the MPFC seed, and found the top-down modulation of consonant/dissonant conditions in non-musicians.

Thirdly, the MVPA searchlight analyses were separately done for the musicians (on frequency interval) and non-musicians (on frequency difference). For musician’s, the lateral STG, medial frontal gyrus, PCC, precuneus, and middle frontal gyrus were mapped for the “perfect fifth vs. tritone” classifications. These midline anatomical structures: including the medial frontal gyrus, PCC, and preneuenus; may suggest that, besides the MPFC and ACC (also along the midline) which were separately revealed by the univariate GLM contrast between the “frequency interval” manipulations, different neural substrates were involved and dug out by different mapping procedures (aka., multivariate mapping). In addition, non-musician’s data revealed that high classification accuracies between roughness and non-roughness manipulations were found in the primary auditory cortex and supramarginal gyrus, suggesting heavier reliance on bottom-up processing in discerning consonant/dissonant tones in non-musicians (see Fig. 4c for details).

As the results of beta-based analyses, which included GLM contrasts, functional connectivities (aka., psycho-physiological interactions), and MVPA searchlight mappings, were described, we now turn to the next layer of beta-based analyses, which incorporated ROI and connectivity-based RSA mappings, and the final spatio-temporal RSA searchlight mappings.

### Representational Similarity Analysis (RSA) results

#### Voxel-wise and network-based RSA results

In this RSA-derived analyses session, 8 different RDMs were constructed based on different ways to encode the relative distances among the 20 trials, according to the 8 RDM models (see Fig. 5, also detailed in the method section, pp. 8). First, the voxelwise trial-estimated beta were compared to the designated RDMs, and the brain regions significantly correlated with each RDM were overlaid, in orange-to-yellow colors (with the corresponding t-values underneath), within the four-view brain surfaces in the top two rows of Fig. 6. For the network-based RSA, the nodes and the edges/sticks within each brain surface were the significant ROIs that were selected from correlation with all-separate RDM for both groups (Fig 6. top row), with the freq-interval RDM for musicians (Fig 6. middle row, left) and with the freq-difference RDM for non-musicians (Fig 6. middle row, right).

**Fig. 5.**
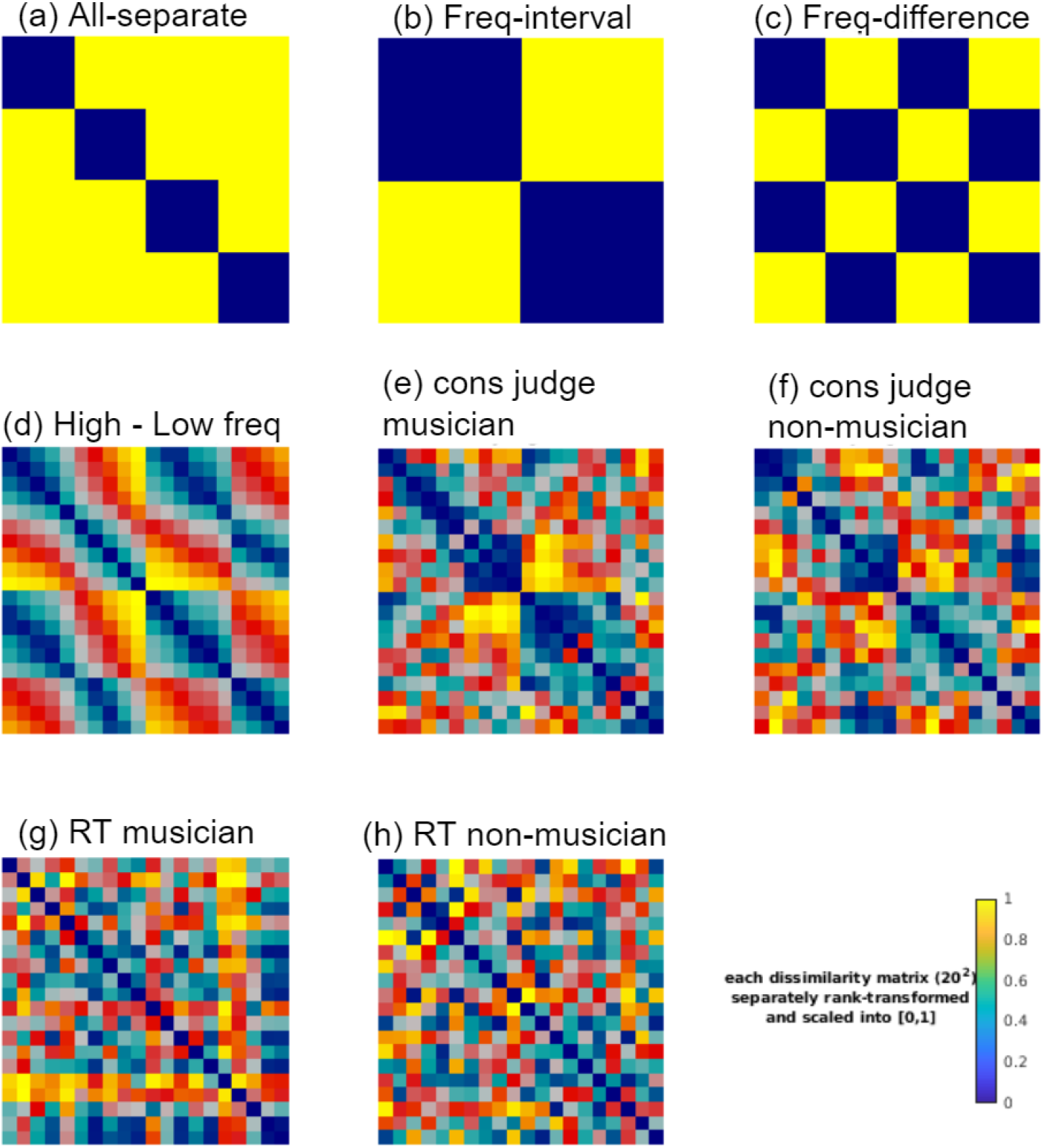
Eight RDM models, representing four different conditions: (a) allSeparate: all 4 (2 perfect fifth/tritone x 2 rough/non-rough) conditions to be equally distant; (b) freq interval; perfect fifth vs. tritione; (c) freq difference: rough vs. non-rough; (d) high and low frequency differences between the two notes (abbreviated as “high-low freq”); (e) average responses of consonant judgments in musicians; (f) average responses of consonant judgments in non-musicians, (g) response times (RTs) of the consonant judgments for musicians; and (h) response times of the consonant judgments for non-musicians.

**Fig. 6.**
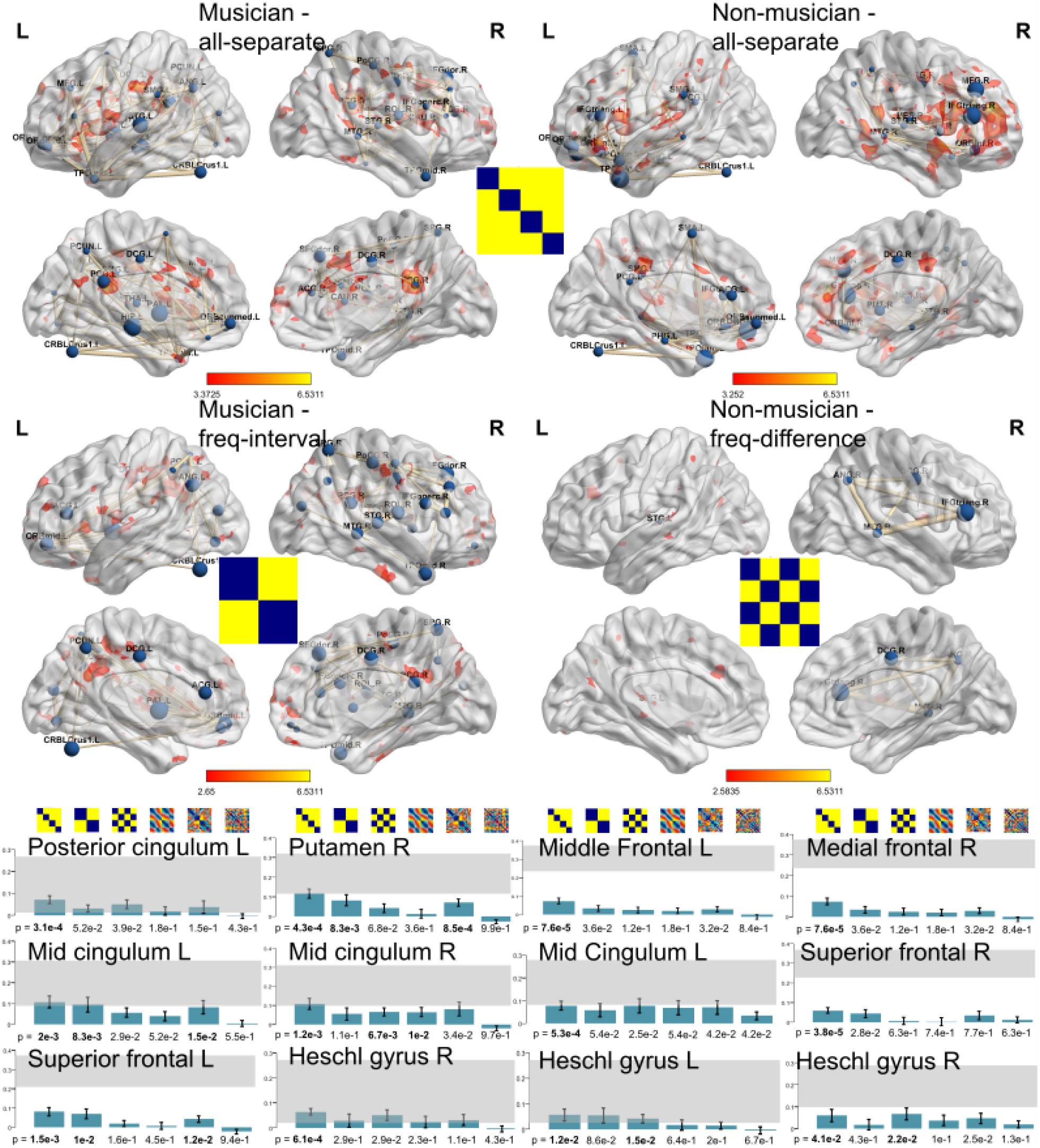
The results of voxel- and ROI/connectivity-RSA. The top row shows the brain areas correlated with all-separate RDM (Representational Dissimilarity Matrix) for musicians (left) and non-musicians (right). The voxel-wise correlations are represented in orange-to-yellow colors, indicating significant correlated regions. The nodes (different sized ‘balls’) and edges (‘sticks’ of various lengths and thicknesses) further show the connectivities among significant ROIs (usually 10 out of 90 cerebral ones). The mid row shows musicians’ brain map correlating with frequency interval RDMs (left) and nonmusicians’ with frequency difference RDMs (right). In the bottom row, the bar graphs represent the brain regions and its relatedness to the 6 model RDMs for each group (left panel for musicians, and right panel non-musicians). The bar is measured by a Spearman correlation between ROI RDM and model RDM in group averages; and the gray area on top shows the ‘noise ceilings’, indicating the maximum performance given the level of noise in the data. The significance *p*-values were under each bar graph, respectively.

As shown in top row of Fig. 6, for musicians (left column) and nonmusicians (right column), both the voxelwise- and ROI connectivity-wise RSA searchlight mappings included the midline areas: bilateral mid- and posterior- cingulate cortex, as well as lateralized auditory-processing areas, but more or less similar (but different emphasis) across both groups. For the middle row, the frequency interval sensitive regions for musicians (left) were again along the midline and lateralized area (including left postcentral gyrus, right superior frontal gyrus, and left insula), among the nodes found significant with freq-interval RDM including the left precuneus, angular gyrus and mid/anterior cingulum cortex are convergely connected to MPFC. For the non-musicians (right), the correlated regions for discerning frequency difference include: the left superior temporal gyrus as voxel-based, and ROI-based RSA showed connected edges among the inferior frontal gyrus, the medial temporal, and the angular gyrus, all in the right hemisphere. The bottom row showed the common 6 ROIs across the top and middle row RSA results: one can see that out of the six *p* values, corresponding to the match with the 6 model RDMs (4+2 for musicians, shown at the bottom row), the all-4-different model always, if not all, got the smallest p values (or best fit against the null models) in 6 chosen listed ROIs, suggesting that the all-different models captured all the condition differences (since all-different means that each of the 2×2 conditions: perfect-fifth rough, perfect fifth non-rough, tritone rough, and tritone non-rough, are all unique in its own right). Besides the above mentioned RDMs, musicians showed right middle temporal gyrus and posterior cingulate regions highly correlated with frequency-difference RDM,and cingulate gyrus for the 4th (high-low freq) model, but none of brain areas were significant for the RT model. Meanwhile, the non-musicians showed highly correlated brain regions in right middle temporal gyrus, left superior temporal gyrus, right cingulate gyrus, and left precuneus with freq-interval model; and bilateral superior temporal gyrus for consonant accuracy (5th model) RDM, and posterior cingulate for the RT (6th) behavior model, but none for the high-low frequency difference (4th) RDM.

#### ssRSA searchlight mapping

Lastly, to fully characterize the spatiotemporal dynamics of participants’ consonance perception by incorporating both the temporal resolution of ERP and the spatial resolution of fMRI, the spatio-temporal RSA was adopted with the temporal RDM (ERP amplitudes over the 20 trials from 8 channels of interest), constructed every 3 ms time bins from 0 to 500ms, correlating with the spatial RDM (fMRI betas over the same 20 trials), constructed over 100 neighboring voxels’ searchlight. By moving this 3ms temporal window along, the (dis)similarity among these 20 trial-related ERP amplitudes (moving) and the fMRI estimated betas (fixed) were correlated to reveal the highly significant brain areas in different time bins. As shown in Fig. 7 (and supplementary movies: 1 for musicians and 2 for non-musicians): as early as 100 ms (or even 50ms) after the stimulus onset, auditory cortex was highly responsive (negatively correlated), for both musicians and nonmusicians. For musicians, the N1-related brain activities were near mPFC (both 117 and 144ms) and r-DLPFC, among other regions; and then spread toward para-mPFC regions (e.g. superior frontal or dorsal anterior cingulate cortex) during the P2 effect (both positively, 230 and 261 ms); In contrast, nonmusicians show N1-related effects with para-mPFC and bilateral superior temporal, and limbic areas (~101ms), and then switched to thalamus (136~230ms), finally to the orbitofrontal and occipital areas (for P2 effects, positively). Around N2 (350ms) latency, dorsolateral prefrontal and dACC were additionally observed only in musicians. All these movies speak to the different spatio-temporal characteristics behind the two populations’ consonance perception dynamics.

**Fig. 7.**
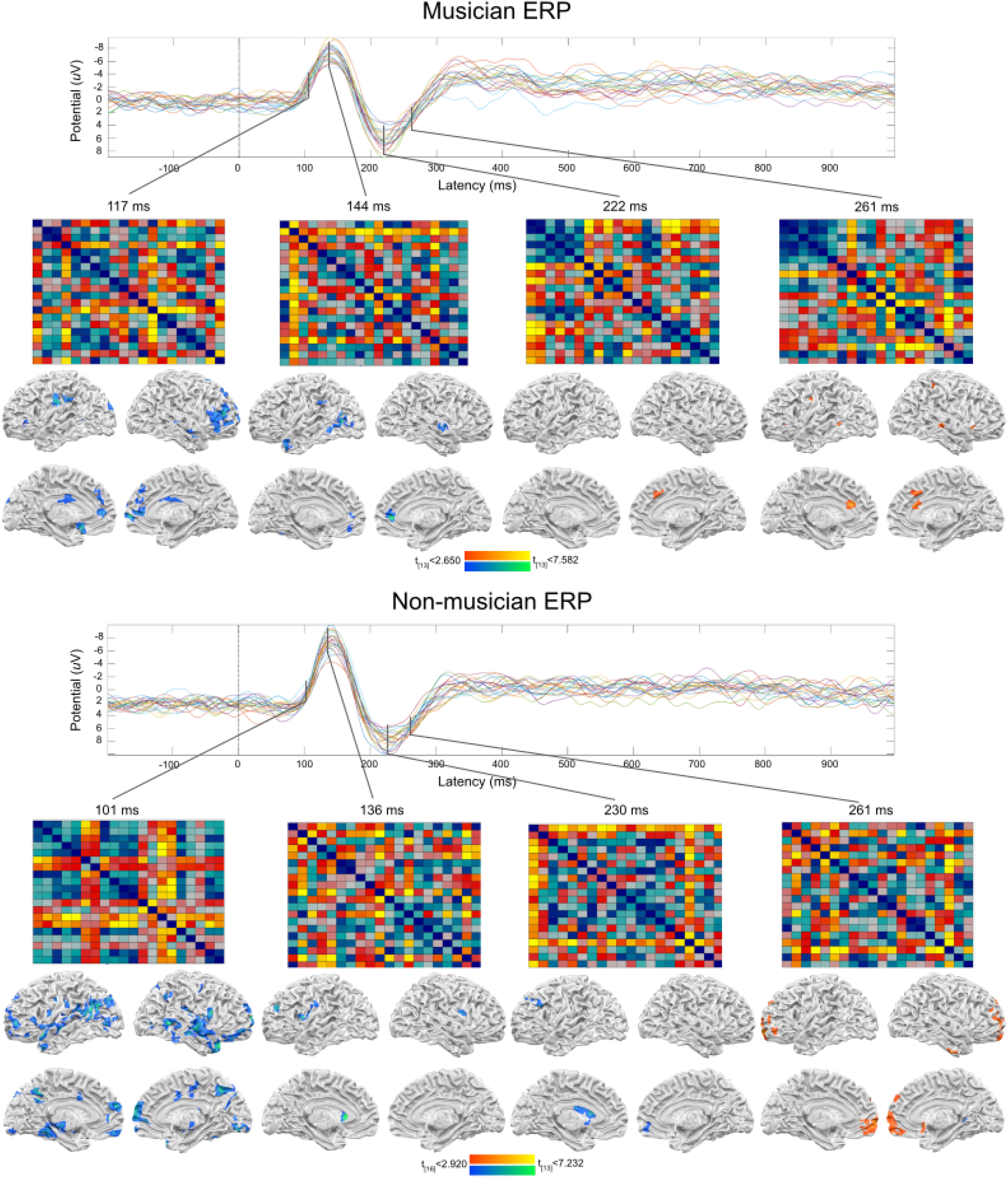
The results of Spatio-temporal Searchlight RSA (ssRSA), according to the temporal order from 0 to 500 ms post-stimuli. The ERP plots show 20 stimulus ERP activations averaged across the 8 channels (including F3, Fz, F4, FC3, FCz, FC4, Cz, and CPz). The ERP RDM generated from 20 ERP activations were further correlated with fMRI voxel RDM group results, thresholded at *p* < .01 (and the cluster threshold 30 voxels) for the brain mappings. Overall, both movies show that (1) both musicians and nonmusicians exhibited auditory negative peaks around 100ms at the auditory cortex, with nonmusicians showed more negatively correlated spots; (2) for musicians, the N1-related brain activities were near mPFC (both 117 and 144ms) and r-DLPFC, among other regions, and then spread toward nearby regions (superior frontal or dorsal anterior cingulate cortex) during the P2 effect (both positively, 230 and 261 ms); (3) In contrast, nonmusicians show N1-related effects with peri-mPFC and bilateral superior temporal, and limbic areas (~101ms), then switched to thalamus (136~230ms), and then finally orbitofrontal and occipital areas (for P2 effects, positively). See the supplementary video 1 and 2 for full movies (between 0 to 500ms), or click the link below. Musicians: https://drive.google.com/file/d/1viug_Tam9Z3x8vOwvCLVjvkd_oxZXjNQ/view?usp=sharing Non-musicians: https://drive.google.com/file/d/12gd4yF6ZVrU67cLdYsZPgBgUweKGENip/view?usp=sharing

## Discussion

The main focus of the study is to investigate the different mechanisms between non-musicians’ and musicians’ brains, and how training induced this change, presumably from the former to the latter. In addition, part of the motivation was to revisit the old finding (Kung et al., 2014) with the extension of fMRI using the same paradigm, with necessary modifications (such as sparse sampling). The behaviour results replicated well with those from the earlier study (c.f., Fig. 4 of Kung et al., 2014 pp. 975, with our current Fig. 2), reassuring the reliance of different features between musicians and non-musicians upon discerning disso-/conso-nance. Second, at least the ERP results in terms of frequency interval (“perfect fifth vs. tritone) in the N1/P2 periods across groups were consistent with those in (Kung et al., 2014; Liang et al., 2016), reinforcing the notion of neural plasticity by top-down influence in musicians (Bailes et al., 2015; Regnault et al., 2001). In contrast, both musicians and non-musicians’ shared modulation of N1/P2 components, along the rough vs. non-rough (aka. frequency difference) dimensions and also along the midline channels, were different from the “group x frequency differences” interaction effects reported in Kung et al., (2014), who showed only non-musicians with higher rough vs. non-rough amplitude differences along the P2 and the midline. To clarify, we reexamined the method details between the Kung et al. (2014) original and the current replication study. Although the two adopted the exact same stimuli and same experimental setup, the two samples of amateur musicians and non-musicians actually composed of different majors and inclusion criteria for musicians: 80% accuracy in Kung et al. (2014), and 75%, or 9 out, of the 12 musicians were pianists (and 1 violinist). In contrast, the current study adopted a slightly lenient (75% accuracy) inclusion criteria, and only 50%, or 7~8 out, of the 15 musicians were pianists (and 5 violinists). Even though three of the four hypothesis sets: the significant effect of frequency interval on musicians, the significant effect of frequency differences on nonmusicians, and the null effect of frequency interval on nonmusicians, are all well replicated across these two samples, the ERP discrepancy for musicians of different majors (e.g., pianists vs. violinists) (Coro et al., 2019) on frequency difference deserves more examinations/replications in the future.

Using the sparse sampling methods, the fMRI experimental results showed the complementary recruitments of task-related brain regions: basically, both the ANOVA (Fig. 4a) and within-group GLM analyses suggested the activated regions were both along the midline (e.g., precuneus, cingulate gyrus, medial frontal gyrus, caudate, etc, in both Fig. 4a and 4b) and lateralized brain areas (Zatorre et al., 2007), e.g., left IFG (Fig. 4a), left superior and middle temporal gyrus (in both Fig. 4b and Supp. Table 2). These findings were in line with one recent meta-analysis study suggesting the involvement of midline brain regions (e.g., ACC), as well as the increases in both the structural and functional connectivities between lateralized auditory and motor cortex with years of longitudinal musicianship (Olszewska et al., 2021). For both musicians (on frequency ratio) and non-musicians (on frequency differences), the rACC/MPFC were consistently engaged in the dissonant processing (again, Fig. 4b and Supp. Table 2). As one recent study linked rACC/MPFC for dissonance with emotional inharmonic information (Bravo et al., 2020), the same area in our study might also be related to the emotional responses to the unpleasantness associated with dissonant chords (Blood et al., 1999). With regard to the multivariate analysis approach, our similar midline plus bi-lateralized findings (Fig. 4c) once again also in agreement with the findings that separated musicians vs. nonmusicians when processing dynamic musical features (Saari et al., 2018), including frontal gyrus, anterior cingulate gyrus (ACG) and right STG. That said, the task (discriminating tonal features vs. consonance judgments) and classification schemes (classifying subjects vs. conditions) differences across studies, among others, might still be considered when comparing different patterns of multivariate analysis results.

The RSA analyses, encompassing the voxel- and connectivity-based renderings on musicians (for frequency ratio) and non-musicians (for frequency differences) (Fig. 6), illuminates more from the network perspective: overall, musicians recruit bilateral midline and lateralized brain networks (mostly around mid-to-superior temporal, inferior frontal, and posterior parietal/precuneus areas) by frequency ratio model; whereas non-musicians more heavily along the orbitofrontal, the rostral ACC network by freq-difference model, when both groups were able to be captured by the all-different models. While there is currently no available publication documenting the use of Representational (Dis-)Similarity Analysis in consonant judgments on musicians or non-musicians, there are 1 case report of a professional singer/composer (Levitin & Grafton, 2016) and another preprint on music listening intervention with healthy older adults (Quinci et al., 2021). As a useful analysis toolkit, RSA models (Fig. 5) and results (Fig. 6) in the current study showcases how the various stimulus features or response properties could be incorporated into the data analysis stream, providing additional insights from different analysis angles.

The last piece contributed by the RSA, the ssRSA searchlight mapping (Salmela et al., 2018), leverages the relative strengths of ERP (temporal) and fMRI (spatial) resolutions into musician/nonmusicians’ brain, by the additional constraints provided by the 20-stimulus orderings (shown on top of Fig. 7 and the 2 supplementary movies, averaged across 8 midline ERP channels). These spatial-temporal couplings between averaged ERP amplitudes and fMRI beta RDMs nicely showed the spatial distributions, for both musicians and nonmusicians, the N1(negatively)-associated brain areas: auditory cortex (50-100ms), the mPFC (144ms); for the P2 component (200~300 ms): for musicians: dorsal anterior cingulate cortex (dACC), superior frontal gyrus. For non-musicians, P2: thalamus (negative), orbitofrontal and occipital (positively correlated). For the N2 stage (350ms), musicians showed again dACC and dorsolateral prefrontal areas. These time-locked brain mappings provide unique brain dynamics into the current stream of results, representing final joint constraints in mapping brain networks in a ~100ms-level resolution. Even though nowadays people push for improvements by simultaneous fMRI-EEG recording (Ritter & Villringer, 2006), the current ssRSA methodology still has its merits, providing additional constraints from the data analysis support of convergence. Together, the human brain mapping of various populations under different task contexts could be more readily and ecologically collected and analyzed.

One obvious benefit yielded by multiple analyses out of a single dataset is the wealth of perspectives illuminated by different angles: diverse findings happen, while converging results provide more robust and compelling evidence (Davis et al., 2011). In the current study, the medial prefrontal cortex (mPFC) was constrained by 4 types of joint analyses: GLM contrasts (Fig. 4a), functional connectivity (Fig. 4b), network-based RSA (Fig. 6, middle left), and finally the ssRSA searchlight mapping (Fig 7, top row, 117~144ms, blue spots), as one of the core networks governing musicians’ top-down consonant perception. Such strong converging evidence is extra implicative when combined with the literature documenting mPFC’s importance in memory and decision making (Euston et al., 2012), social perception (Isoda, 2021), and music cognition (Bravo et al., 2020; Mizuno & Sugishita, 2007), especially from the frequency ratio perspective. In addition, lateralized superior temporal cortex (STG) or auditory cortex, constrained by GLM contrasts, functional connectivity, MVPA, and the ssRSA (~50-100ms), were not surprisingly also among one of the earlier consonance perception networks for musicians as well. For the P2 component (250-350ms), musicians’ superior frontal cortex and dACC were also implicated in processing musical features: such as timbre, rhythm, and tonality (Saari et al., 2018), and imagined music performance (Tanaka & Kirino, 2019). Taken together, the midline regions (here mPFC and dACC) and lateralized brain areas (e.g., auditory cortex or STG) are responsible for musicians’ top-down and/or integrated processing consonant perception. For nonmusicians, the story is still more rough vs. non-nough oriented: for example, the early recruitment of mPFC (by ssRSA around ~100ms), concurred by GLM contrast (along the rough vs. non-rough dimension). Besides, these stimulus-correlated RSA areas lasted longer (into the P2, <250ms, stage) than the musician counterparts, and included the negatively correlated thalamus region (Joris et al., 2004) into the presumably P2 component (150-250ms window), reflecting nonmusicians’ longer dependence on bottom-up processing for tonal features (e.g., roughness).

In sum, the current study tries to capture the spatio-temporal dynamics underlying the recruitments of midline and lateralized brain areas, revealed by both ERP and fMRI and the associated separate and joint analyses, at different time windows and for musicians and nonmusicians, respectively. It not only replicates our previous ERP findings mostly (Kung et al., 2014), but also elucidate more on the target brain areas associated with each ERP component (and for different subject groups). The current study, along with similar endeavors (Crespo-Bojorque et al., 2018), each provides their own vantage point, could join the wave to move closer into the neural underpinnings of the targeted behavior, here the different processing characteristics underlying consonant vs. dissonant judgments.

## Acknowledgments

the authors like to thank the support of the Ministry of Science and Technology (MOST 103-2410-H-006-030), the Humanistic-Social Benchmarking Project from the Ministry Of Education, as well as the consultation and instrument availability by the Mind Research and Imaging (MRI) Center at NCKU.

## Notes

Conflict of interest statement: The authors declare no competing conflict of interest.

### Competing Interest Statement

The authors have declared no competing interest.

