## Supplementary materials for "Musicians and non-musicians’ consonant/dissonant perception investigated by EEG and fMRI"

Author Affiliations:

Contents:

Supplementary Figure 1. fMRI GLM-ANOVA results in brain map

Supplementary Table 1. Cluster-table of fMRI ANOVA results

Supplementary movie 1. Spatio-Searchlight RSA results for musician

Supplementary movie 2. Spatio-Searchlight RSA results for non-musician

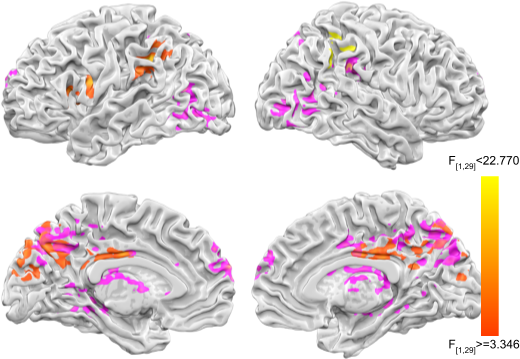

Supplementary Figure 1. GLM ANOVA results, main group effects color coded in orange, pink color indicate interaction effect between group X frequency difference (rough and non-rough condition), and the yellow patch represent interaction effect between group X frequency ratio (tritone and perfect fifth condition), all significant areas are above the *p* < .05 with the FWE corrected by alphasim.

Main effect of group

-----------------------------------------------------------

x	y	z |	k |	max	|	mean	| tdclient

-10 -12	24 |	455 | 22.734737 | 6.401255 |LH Caudate (Caudate Body) (-12; -12; 21) [d=3.6mm]

-42	14	19 |	192 | 22.357069 | 7.153583 |LH Inferior Frontal Gyrus (Brodmann area 9) (-45; 16; 22) [d=4.7mm]

-47 -42	32 |	201 | 20.195290 | 7.069099 |LH Supramarginal Gyrus (Brodmann area 40) (-44; -42; 34) [d=3.6mm]

Main effect of within rough/non-roughness conditions

-----------------------------------------------------------

x	y	z |	k |	max	|	mean	| tdclient

-61 -19	8 |	222 | 21.628088 | 6.620202 |LH Transverse Temporal Gyrus (Brodmann area 42) (-61; -19; 10) [d=2.0mm]

58 -39	7 |	187 | 17.478743 | 6.728539 |RH Middle Temporal Gyrus (Brodmann area 22) ( 58; -39; 5) [d=2.0mm]

6 -23	8 |	99 | 15.542187 | 6.098053 |RH Thalamus (Pulvinar)

Interaction effect between group and rough/non-roughness conditions

-----------------------------------------------------------

x	y	z |	k |	max	|	mean	| tdclient

-25	-5	13 |	714 | 22.866367 | 6.577911 |LH Lentiform Nucleus (Putamen)

-4	65	9 |	98 | 21.946060 | 7.207768 |LH Medial Frontal Gyrus (Brodmann area 10)

28 -37	31 |	735 | 20.537809 | 6.530092 |RH Cingulate Gyrus (Brodmann area 31) ( 21; -40; 31) [d=7.6mm]

42 -77	6 |	154 | 17.659258 | 6.527731 |RH Middle Occipital Gyrus (Brodmann area 19)

31 -16	-3 |	97 | 16.366350 | 6.721217 |RH Lentiform Nucleus (Putamen)

-21 -24	-4 |	451 | 15.946597 | 6.448032 |LH Sub-lobarLateral Geniculum Body (-21; -24; -2) [d=2.0mm]

-18	29	14 |	97 | 14.352325 | 6.578186 |LH Anterior Cingulate (Brodmann area 32) (-17; 32; 18) [d=5.1mm]

53 -36	25 |	96 | 12.906526 | 6.882773 |RH Inferior Parietal Lobule (Brodmann area 40) ( 54; -34; 26) [d=2.4mm]

Main effect of within tritone/perfect fifth conditions

-----------------------------------------------------------

x	y	z |	k |	max	|	mean	| tdclient

16	8	-2 |	504 | 17.637510 | 6.926910 |RH Lentiform Nucleus (Putamen)

24 -29	3 |	132 | 15.725543 | 6.825944 |RH Thalamus (Pulvinar)

-22 -69	-6 |	200 | 9.647047 | 5.731200 |LH Lingual Gyrus (Brodmann area 19)

Interaction effect between group and tritone/perfect fifth conditions

-----------------------------------------------------------

x	y	z |	k |	max	|	mean	| tdclient

40 -44	26 |	91 | 10.665401 | 5.527355 |RH Insula (Brodmann area 13) ( 41; -44; 21) [d=5.1mm]

Supplementary Table 2. Cluster table of fMRI ANOVA results, results are shown F-map with degrees of freedom in 1,29 at *p* < 0.05 with peak coordinates.

Supplementary Movie 1. The musician’s ssRSA created from 0 to 500ms ERP from 8 channels (F3, Fz, F4, FC3, FCz, FC4, Cz, and CPz) for every 3ms time bins and its corresponding brain regions based on fMRI data. The video has been uploaded and shared in the following link: <https://drive.google.com/file/d/1viug_Tam9Z3x8vOwvCLVjvkd_oxZXjNQ/view?usp=sharing>

Supplementary Movie 2. The non-musician’s ssRSA created from 0 to 500ms ERP from 8 channels (F3, Fz, F4, FC3, FCz, FC4, Cz, and CPz) for every 3ms time bins and its corresponding brain regions based on fMRI data. The video has been uploaded and shared in the following link:

<https://drive.google.com/file/d/12gd4yF6ZVrU67cLdYsZPgBgUweKGENip/view?usp=sharing>
